# PIK3CA^H1047R^-induced paradoxical ERK activation results in resistance to BRAF^V600E^ specific inhibitors in BRAF^V600E^ PIK3CA^H1047R^ double mutant thyroid tumors

**DOI:** 10.1101/164533

**Authors:** Matthias A. Roelli, Dorothée Ruffieux-Daidié, Amandine Stooss, Oussama ElMokh, Wayne A. Phillips, Matthias S. Dettmer, Roch-Philippe Charles

**Affiliations:** Institut für Biochemie und Molekulare Medizin, Universität Bern, Bern, Switzerland; Institut für Pathologie, Universität Bern, Bern, Switzerland; Cancer Biology Laboratory, Peter MacCallum Cancer Centre, Melbourne, Australia

**Keywords:** Aggressive thyroid cancer, BRAF^V600E^ inhibitor resistance, PI3’K inhibitors, paradoxical ERK activation, overcoming resistance

## Abstract

Thyroid carcinomas are the most prevalent endocrine cancers. The BRAF^V600E^ mutation is found in 40% of the papillary type and 25% of the anaplastic type. BRAF^V600E^ inhibitors have shown great success in melanoma but, they have been, to date, less successful in thyroid cancer. About 50% of anaplastic thyroid carcinomas present mutations/amplification of the phosphatidylinositol 3’ kinase. Here we propose to investigate if the hyper activation of that pathway could influence the response to BRAF^V600E^ specific inhibitors.

To test this, we used two mouse models of thyroid cancer. Single mutant (BRAF^V600E^) mice responded to BRAF^V600E^-specific inhibition (PLX-4720), while double mutant mice (BRAF^V600E^; PIK3CA^H1047R^) showed resistance and even signs of aggravation. This resistance was abrogated by combination with a phosphoinositide 3-kinase inhibitor. At the molecular level, we could show that this resistance was concomitant to a paradoxical activation of the MAP-Kinase pathway, which could be overturned by phosphoinositide 3-kinase inhibition *in vivo* in our mouse model and *in vitro* in human double mutant cell lines.

In conclusion, we reveal a phosphoinositide 3-kinase driven, paradoxical MAP-Kinase pathway activation as mechanism for resistance to BRAF^V600E^ specific inhibitors in a clinically relevant mouse model of thyroid cancer.

## Introduction

Thyroid cancer is the most frequent form of endocrine malignancy. <u>P</u>apillary <u>t</u>hyroid <u>c</u>ancer (PTC) is the most prevalent type of thyroid carcinoma (80%). It is moderately aggressive, moderately lymphometastatic and has a response rate above 90% to standard radioiodine treatment. The main risk is a possible progression (5-10% of the cases) to more aggressive variants including radio-iodine-resistant PTC, poorly differentiated thyroid carcinoma and <u>a</u>naplastic <u>t</u>hyroid <u>c</u>arcinoma (ATC) [1]. For ATC, the survival rate is very poor with a median survival of 2-7 months after diagnosis and a mortality rate of 90% within the first year after diagnosis [2]. Metastases can be observed in 10-20% of ATC cases, mainly in the lung and bones [3]. However, unlike most cancer patients, ATC patients do not predominantly succumb to “generalization” of the disease, but rather to local invasion of the tumor into the tracheal space. Undifferentiated cancer cells invade the space between the cartilage ring and the tracheal epithelium inducing dyspnea and suffocation [4]. The thyroid’s localization next to major vessels like the carotids makes complete curative resection impossible. There is currently no efficient treatment for ATC, the disease being highly resistant to radio-and chemotherapies, including radioiodine I^131^ that usually yields excellent therapeutic outcome in other thyroid cancers.

BRAF is part of the canonical signaling pathway RAS→RAF→MEK→ERK hereafter termed MAPK pathway. The mutant gene coding for BRAF^V600E^ is found in almost 45% of PTCs and 20-40% of ATCs, making it the most common genetic alteration in thyroid cancer (˜40% overall) [5,6]. Independent models expressing the BRAF^V600E^ mutation specifically in the thyroid demonstrate that this constitutively active form of the protein leads to development of PTC [7,8], thus confirming the importance of the mutation in pathogenesis. More recent models show that BRAF^V600E^ can collaborate with p53 deletion [9] or with PIK3CA^H1047R^ mutation [10] to promote PTC progression to ATC.

The phosphatidylinositol 3’ kinase or PI3’K is part of the RAS→PI3’K→AKT→mTOR pathway hereafter termed PI3’K pathway. PI3’K phosphorylates phosphoinositides on the 3 position of the inositol ring [11] resulting in AKT recruitment to the membrane and its phosphorylation [12]. There are several PI3’K which are heterodimers composed of a catalytic and a regulatory subunit. *PIK3CA* codes for the p110α catalytic subunit of class I PI3’K [13]. The PIK3CA^H1047R^ mutation renders the protein constitutively active and can be found frequently in cancers [14]. PI3’K signaling alterations frequently occur in aggressive thyroid cancers with 40% gene amplification and 20% mutations [15,16].

Pharmacological mutation specific inhibition of BRAF^V600E^ in melanoma patients with vemurafenib leads to a dramatic tumor regression [17–19]. Unfortunately, half of the patients relapse after six months of treatment. Many routes to acquired resistance have been proposed, including elevated expression of CRAF [20] or BRAF kinases or aberrant expression of a BRAF splice variants [21–23]. All these resistance mechanisms lead to the reactivation of RAF kinases. In addition, several other means of acquired resistance are described involving mutations in other partners of the MAPK pathway such as N-RAS [24] or MEK [25]. Remarkably, all these mechanisms result in MAPK pathway reactivation, demonstrated by ERK phosphorylation, and eventually lead to a resumption of tumor growth. This emphasizes the central role of the MAPK pathway as the main driver of tumor growth and resistance and the necessity to pharmacologically target that pathway to achieve tumor reduction.

Even though 40% of all thyroid tumors harbor the BRAF^V600E^ mutation, unlike in melanoma, it is not clear whether BRAF^V600E^ inhibition could be used against thyroid tumors. About 10% or thyroid cancers are incurable because of their diffuse presentation making them inoperable as well as their loss of iodine hunger. New targeted therapies are therefore urgently needed for ATC and radio-iodine-resistant PTC. There are some encouraging results from single case studies in BRAF^V600E^ positive ATC [26,27] and also for invasive BRAF^V600E^ positive PTC albeit from reports of very small cohorts [28]. The biggest study so far concerns 7 ATC patients with various responses from complete/partial regression but surprisingly also to tumor progression [29]. Overall these studies suggest an approximate 50% response rate to BRAF^V600E^ inhibitors in aggressive thyroid cancer. One case even showed patient’s rapid worsening which is the actual opposite of what was expected after vemurafenib treatment [27]. This suggests “pre-existing” drug resistance that does not result from treatment adaptation over a few weeks like in melanomas.

Understanding these refractory forms of cancer is essential for efficient use of targeted therapies. In this study, we used a BRAF^V600E^ PIK3CA^H1047R^ double mutant, as well as a BRAF^V600E^ single mutant mouse model to investigate the tumor burden response to BRAF^V600E^ specific inhibition. Our experiments showed that the PIK3CA^H1047R^ mutation conferred resistance to the drug, and this resistance was waived by combination treatment with a PI3’K inhibitor. The resistance was correlated to paradoxical hyperactivation of ERK, that was also depending on PI3’K activity.

## Results

### BRAF^V600E^ single mutant thyroid tumor respond to PLX4720 inhibition, while BRAF^V600E^; PIK3CA^H1047R^ double mutant tumors do not

Our primary aim was assessing the effect of BRAF^V600E^ inhibition in our two mouse models. The BRAF^V600E^ single, and the BRAF^V600E^/PIK3CA^H1047R^ double mutant. PLX-4720, a commonly used pre-clinical surrogate for vemurafenib [30–32] that is similarly potent but more soluble/bioavailable [33] was used to inhibit BRAF^V600E^ specifically.

Mice were bred, tumors induced at the age of one month and the treatments started two months after tumor induction to allow tumors to form (Fig. 1A), but still leave enough time to treat mice for 3 more months without reaching 6 months after induction when mice usually reach endpoint (whistling, breathing issues and consequent sudden weight loss) (Fig 1A).

**Figure 1:**
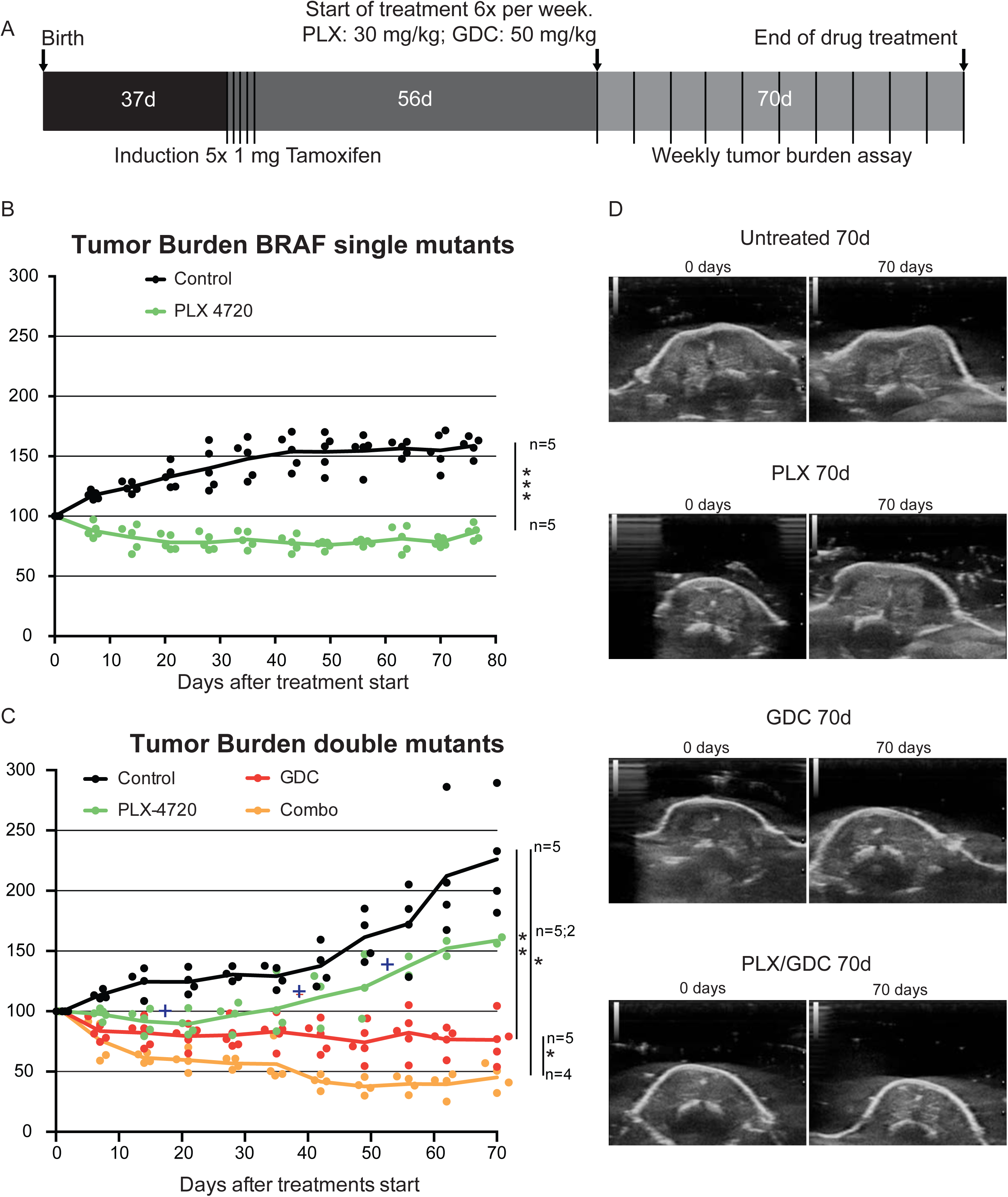
PI3’K inhibition counteracts PLX-4720 resistance in a double mutant thyroid cancer mouse model. **(A)** Schematic timeline depiction of *in vivo* Experiment. Tumor burden in single mutant *Braf*^*CA/+*^;*ThyroglobulinCre*^*ERT2*^ mice **(B)** and double mutant *Braf*^*CA/+*^;*Pik3ca*^*Lat/+*^;*ThyroglobulinCre*^*ERT2*^ mice **(C)**. Crosses represent events of mice reaching end-point criteria. **(D)** Representative pictures from ultrasound imaging of double mutant mice after 70 days of treatment used for assessment of the tumor burden. Thyroid outlines are shown by blue lines.

In single mutant mice, the PLX-4720 drug treatment induced a significant tumor size reduction (-20%) that lasted for more than 10 weeks, while controls increased in size by more than 60% (Fig. 1B). In double mutant mice, tumors were bigger and grew faster than in control mice. PLX-4720 treated animals had a modest drug response for 3 weeks (-10%), then tumor burden resumed growth at a similar rate to control mice with a tumor burden increase of +12 percentage points per week (Fig. 1C&D). Much to our concern, 3 out of 5 animals from this group had to be removed during the course of the experiment as they were reaching predefined endpoint criteria (see above paragraph). This worsening of the condition was only witnessed in PLX-4720 treated animals. GDC-0941 treated animals presented an initial tumor burden reduction of 20% then tumor size stabilized for the rest of the treatment period. Interestingly, when a drug combination of PLX-4720 and GDC-0941 was administered, mice showed a robust response with 60% lower tumor burden after 6 weeks followed by stabilization until the end of the experiment (Fig. 1C&D).

Double mutant tumors present histological worsening after BRAF^V600E^ specific inhibition.

After 10 weeks of treatment, 3 double mutant mice per group were dissected and their thyroids were processed for histological analysis. Control mice had the expected histology: aggressive PTC with tall cell variant, and foci presenting signs of phenotypic progression to ATC (Fig. 2A). GDC-0941-treated mice had smaller thyroid sections, while presenting a similar histology compared to controls (Fig. 2B) with a mixture of PTC containing small ATC foci. Animals treated with the drug combination also presented smaller thyroid sections but in these PTC areas were almost absent, leaving connective tissue and cholesterol clefts, but ATC foci remained (Fig. 2C).

**Figure 2:**
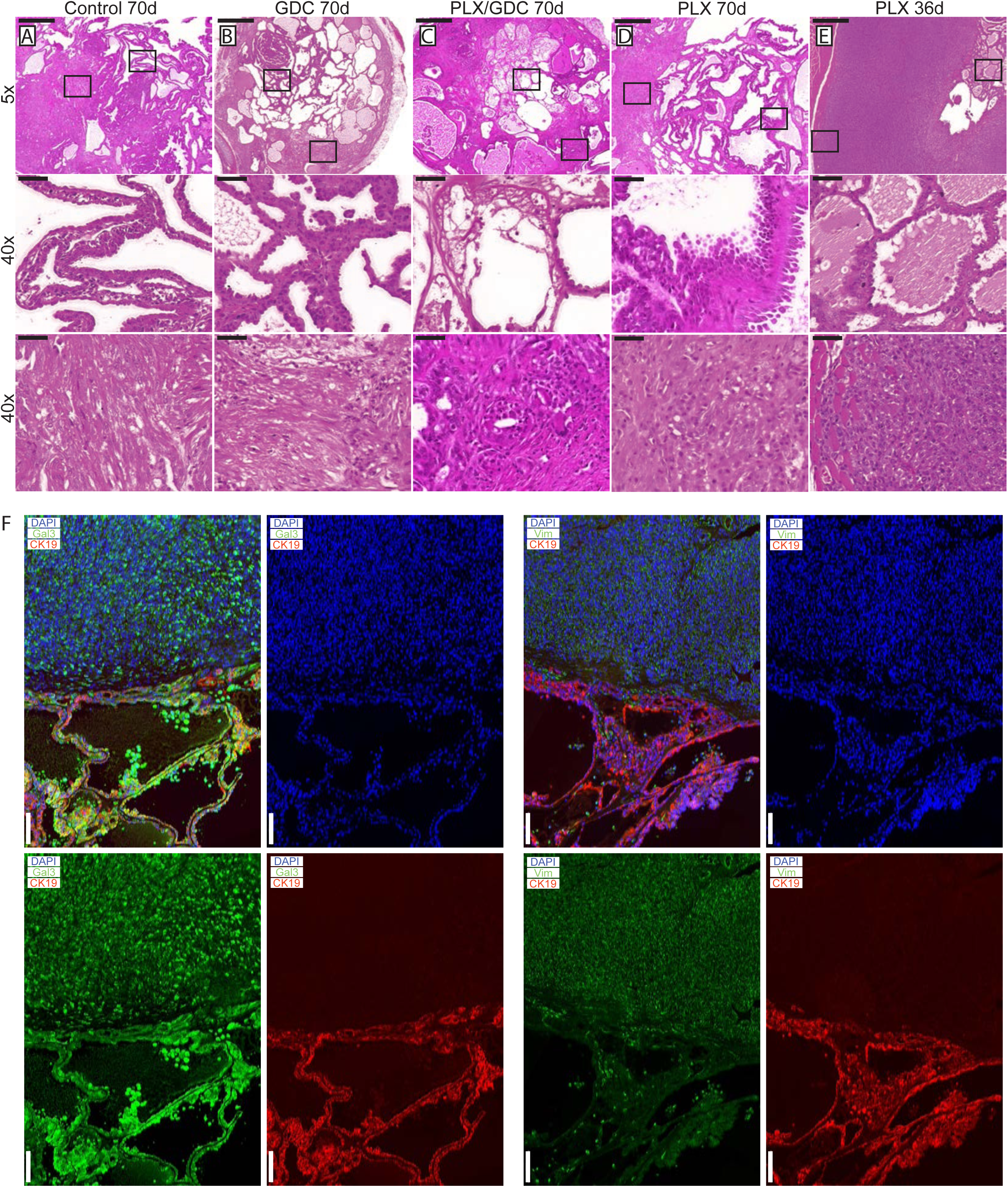
Histological presentation of tumors after 70 days of drug treatments. H&E staining from formalin fixed paraffin embedded and sectioned (5 μm) thyroids from double mutant mice after 70 days of treatment. Representative thyroid tissues at the end of the treatment of **(A)** control, **(B)** 50 mg/kg GDC-0941, **(C)** drug combination (50 mg/kg GDC-0941 and 30 mg/kg PLX-4720 and **(D)** 30 mg/kg PLX-4720 treated mice. (**E)** H&E staining of thyroid tissues from a double mutant mouse treated with PLX-4720 reaching endpoint after 36 days of treatment. Scale bars: 500 μm (upper panels) 50 μm (lower panels). **(F)** Immunostainings of representative PTC and ATC tissues stained for Cytokeratin 19 (red) DAPI/nuclei (blue) and either Galectin-3 or Vimentin (green). Scale bars 100 μm.

PLX-4720-treated thyroids presented large PTC nodule areas that were more solid and featuring areas of hobnail type PTC (Fig. 2D). These sections presented also larger foci that were clear ATC, invading the surroundings (muscles) and eventually the tracheal rings. Overall, they looked more progressed than the controls or any other groups. Finally, early-terminated subjects from the group treated with PLX-4720 presented histology similar to other PLX-4720 treated tumors but some with an even worse histology with one case of full-blown ATC covering about 50% of the section (Fig. 2E). Immunostainings were preformed to confirm the pathological analysis. PTC areas were positive for galectin-3 and cytokeratin 19 while ATC areas were positive for galectin-3 and vimentin (Fig. 2F).

### Double mutant tumors present paradoxical activation of ERK after 10 weeks of PLX-4720 treatment

Signaling pathways in the tumors were investigated by western blot. PLX-4720-treated animals displayed a paradoxical elevated ERK1/2 phosphorylation ratio compared to the controls after 10 weeks of treatment. When treated with GDC-0941 alone, ERK1/2 phosphorylation was unchanged compared to controls. When treated with drug combination, ERK paradoxical activation was abolished resulting in ERK phosphorylation level comparable to controls. AKT phosphorylation was not affected by PLX-4720 while GDC-0941 treatment resulted in a small but significant reduction of AKT phosphorylation (Fig. 3A).

**Figure 3:**
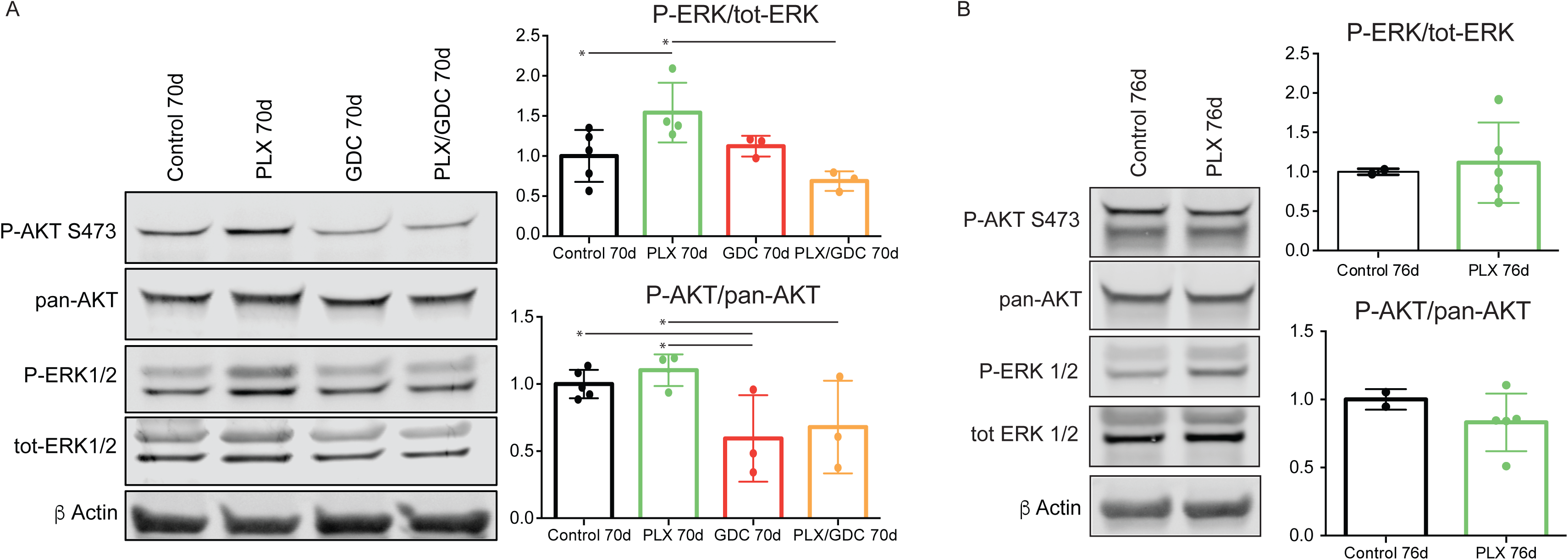
BRAF^V600E^ and PIK3CA^H1047R^ tumors present a paradoxical activation of ERK under PLX-4720 treatment after 10 weeks. **(A)** Representative pooled western blot (left), and quantification of single sample blots (right) of proteins from BRAF^V600E^/PIK3CA^H1047R^ double mutant mice treated for 70 days with either vehicle, PLX-4720 (30 mg/kg), GDC-0941 (50 mg/kg) or a combination of both. **(B)** Representative pooled western blot (left), and quantification of single sample blots (right) of proteins from BRAF^V600E^ single mutant mice treated for 76 days with either vehicle or PLX-4720 (30 mg/kg). Error bars represent standard deviation. Values represent means of calculated ratios. Points represent single ratio values.

For comparison, we performed western blots with proteins extracted from BRAF^V600E^ single mutant mice. In this case, after 70 days of PLX-4720 treatment, ERK phosphorylation in treated animals was comparable to the non-treated. As expected, PLX4720 treatment did not affect AKT phosphorylation significantly (Fig. 3B).

### Paradoxical activation of ERK also occurs after short periods of PLX-4720 treatment

To confirm the observed paradoxical ERK activation, we treated mice with the same drugs for a shorter period of 10 days. We observed the same trend for tumor burden as seen after 10 weeks (Fig. 4A). Only GDC-0941 and drug combination treated animals showed tumor burden reduction. PLX-4720 treated animals displayed an elevation in tumor burden that was not statistically different from the controls but from the two other groups (Fig 4A).

**Figure 4:**
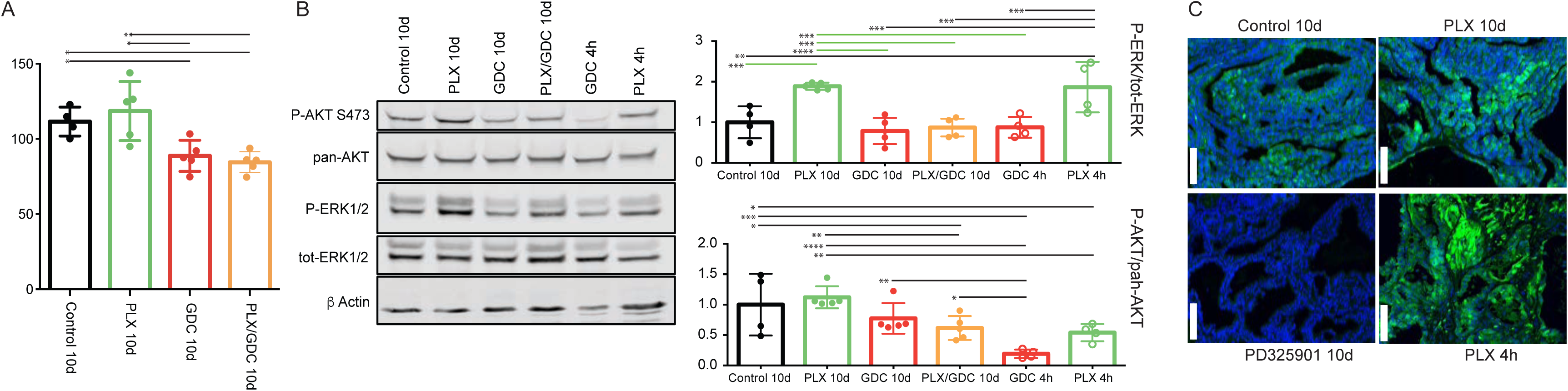
BRAF^V600E^ and PIK3CA^H1047R^ tumors present a paradoxical activation of ERK under PLX-4720 treatment also after 10 days. **(A)** Tumor burden quantification from BRAF^V600E^/PIK3CA^H1047R^ double mutant mice treated for 10 days with either vehicle, PLX-4720 (30 mg/kg), GDC-0941 (50 mg/kg) or a combination of both, and one dose treatments with either PLX-4720 (30mg/kg) or GDC-0941 (50 mg/kg). **(B)** Representative pooled western blot (left), and quantification of single sample blots (right) of proteins from the same animals and from animals treated only once with either PLX-4720 (30 mg/kg), GDC-0941 (50 mg/kg) and dissected 4 hours later. (**C)** Immunofluorescence staining for P-ERK of formalin fixed, paraffin embedded thyroid tumor samples sectioned to 5μm from BRAF^V600E^, PIK3CA^H1047R^ double mutant mice treated with either vehicle, PLX-4720 (30 mg/kg or PD-325901 (5 mg/kg) for 10 days or PLX-4720 (30 mg/kg) for 4 hours. Scale bars: 100 μm.

Similarly to the observed effect at 10 weeks, after 10 days of treatment we observed paradoxical activation of ERK under PLX-4720 treatment that was abrogated by the drug combination (Fig. 4B). To complete this part, we also treated double mutant animals only once with PLX-4720 or GDC-0941 and dissected the animals four hours after drug administration. At this time, the PLX-4720 driven paradoxical activation of ERK was already observable. GDC-0941 treatment on the other hand led to a pronounced de-phosphorylation of AKT.

Finally, to ensure that ERK phosphorylation signal was coming from tumor cells, we performed immunofluorescence staining on tumor samples. The signal was clearly located in the tumor cells and not in the mesenchyme or coming from infiltrating immune cells. The specificity of our staining was verified by treating mice with the potent MEK1/2 inhibitor PD-325901 [34] that completely abrogated ERK phosphorylation after 10 consecutive days of treatment (Fig. 4C).

### Drug combination induces increased cell death in vivo

Searching for the mechanisms driving the observed tumor reduction, we performed immunofluorescence staining for Ki67. We could only detect a significant increase of Ki67 index in PLX-4720 treated animals compared to the drug combination group (Fig. 5A). Then we performed TUNEL staining to monitor DNA fragmentation. The count of apoptotic bodies was increased in GDC-0941 and drug combination treatment (Fig. 5B). In addition, we detected a reduced count in PLX-4720 treated tumors. We then performed a Masson’s trichrome staining to evidence collagen accumulation stained in blue (Fig. 5C). Interestingly, the drug combination treatment led to a significantly increased proportion of collagen rich areas inside the thyroid lobes, evidencing higher tumor regression. Tumors from PLX-4720-treated mice had a significantly lower proportion of collagen in the tumor tissue.

**Figure 5:**
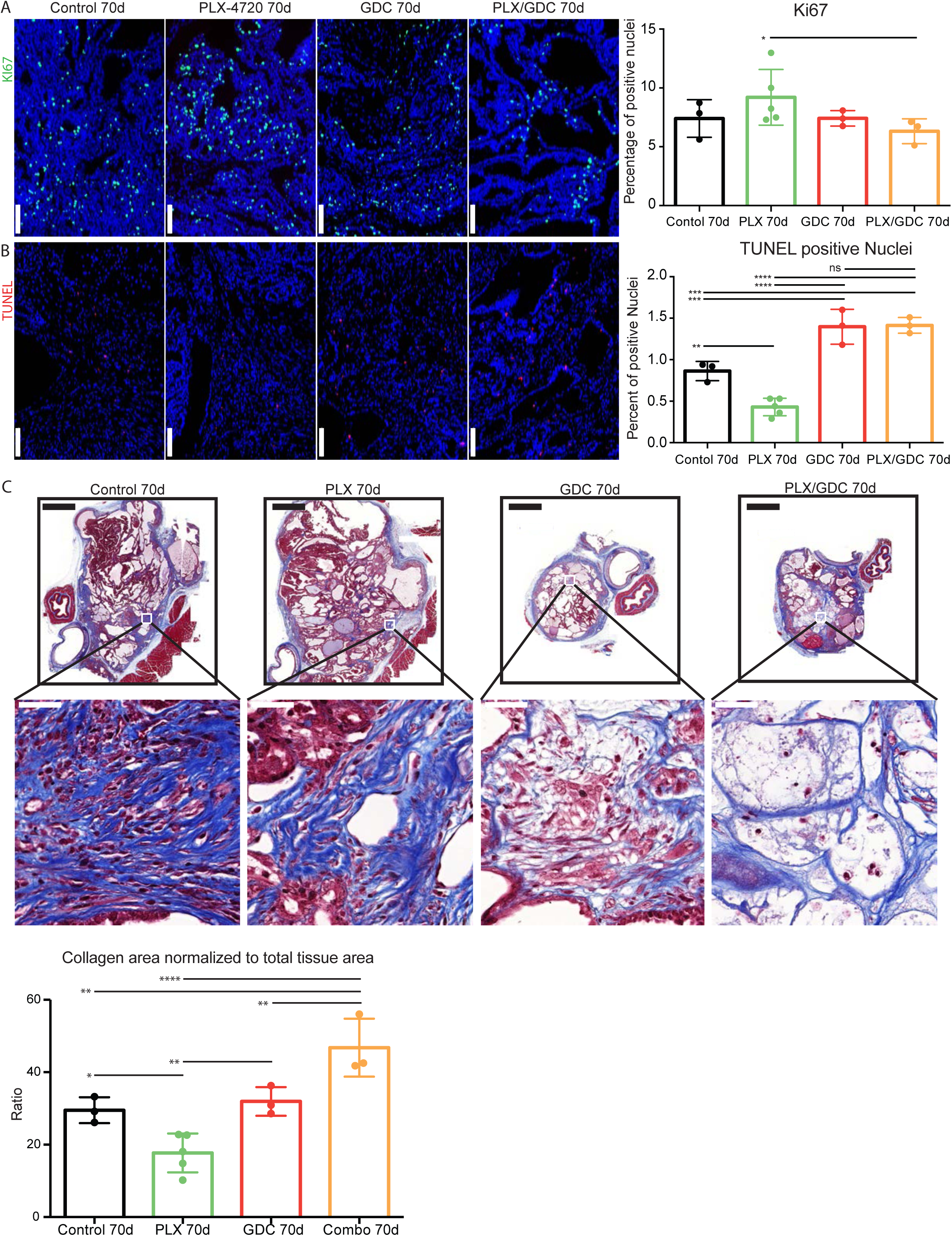
Drug combination induces increased cell death in vivo. Ki67 staining formalin fixed, paraffin embedded 5 μm tumor samples (**A)** Left side: Representative immunofluorescence after 70 days of treatment. DAPI (blue) Ki67 (green). Scale Bars: 100 μm. Right side: Quantification on whole tumors averaged from 4 consecutive sections. (**B)** Left side: TUNEL staining revealed with streptavidin-Alexa633 fluorescent probe (purple) and DAPI (blue). Scale Bars: 100 μm. Right side: TUNEL positive nuclei normalized to total nuclei counts. Scale bars: 100 μm. (**C)** Top: Representative images of Masson’s trichrome stained tumor sections and higher magnifications thereof below. Scale bars: 1 mm (upper) and 50 μm (lower). Bottom: evaluation of collagen rich area normalized to total tissue area per tumor. Error bars represent standard deviation. Values represent means of calculated ratios. Points represent single ratio values.

### Paradoxical activation of ERK under BRAF^V600E^ specific inhibition treatment is also PI3’K-dependent in human ATC cell lines

To further demonstrate that paradoxical activation of ERK also depends on PI3’K signal in human ATC cells, we used three cell lines. OCUT-2 cells harbor the same mutations as the double mutant mouse model (BRAF^V600E^/PIK3CA^H1047R^), whereas 8505c and SW1736, analogous to the single mutant mice, only have the BRAF^V600E^ mutation. The mutational status of *PIK3CA* in these cell lines was confirmed by sequencing [35]. As paradoxical ERK activation has been described to be dependent on drug concentration [36], we exposed the cells to decreasing concentrations of PLX-4032 (vemurafenib). Interestingly, OCUT-2 cells had a greater than 2-fold increase in p-ERK/tot-ERK ratio compared to vehicle treated cells, at low concentrations: 1.6 and 8 nM (Fig. 6A). Paradoxical ERK hyper-phosphorylation was not detectable when cells were subjected to GDC-0941 in addition to PLX-4032 drug dilutions. When we treated them with the same concentrations of PLX-4032, 8505c and SW1736 cells did not exhibit paradoxical ERK activation (Fig. 6B). The additional treatment with GDC-0941 did not affect either the baseline ERK phosphorylation or the response to PLX-4032 regardless of the PLX-4032 concentration in 8505c cells. To strengthen this point, we have also performed a similar experiment using BKM-120, another PI3’K inhibitor (Supp Fig. 1). In this case, BKM-120 similarly abrogated the paradoxical effect seen in OCUT-2, demonstrating that the phenomenon observed is unlikely to be caused by a possible off-target effect of GDC-0941 but was rather specific to its inhibitory effect on PI3’K activity.

**Figure 6:**
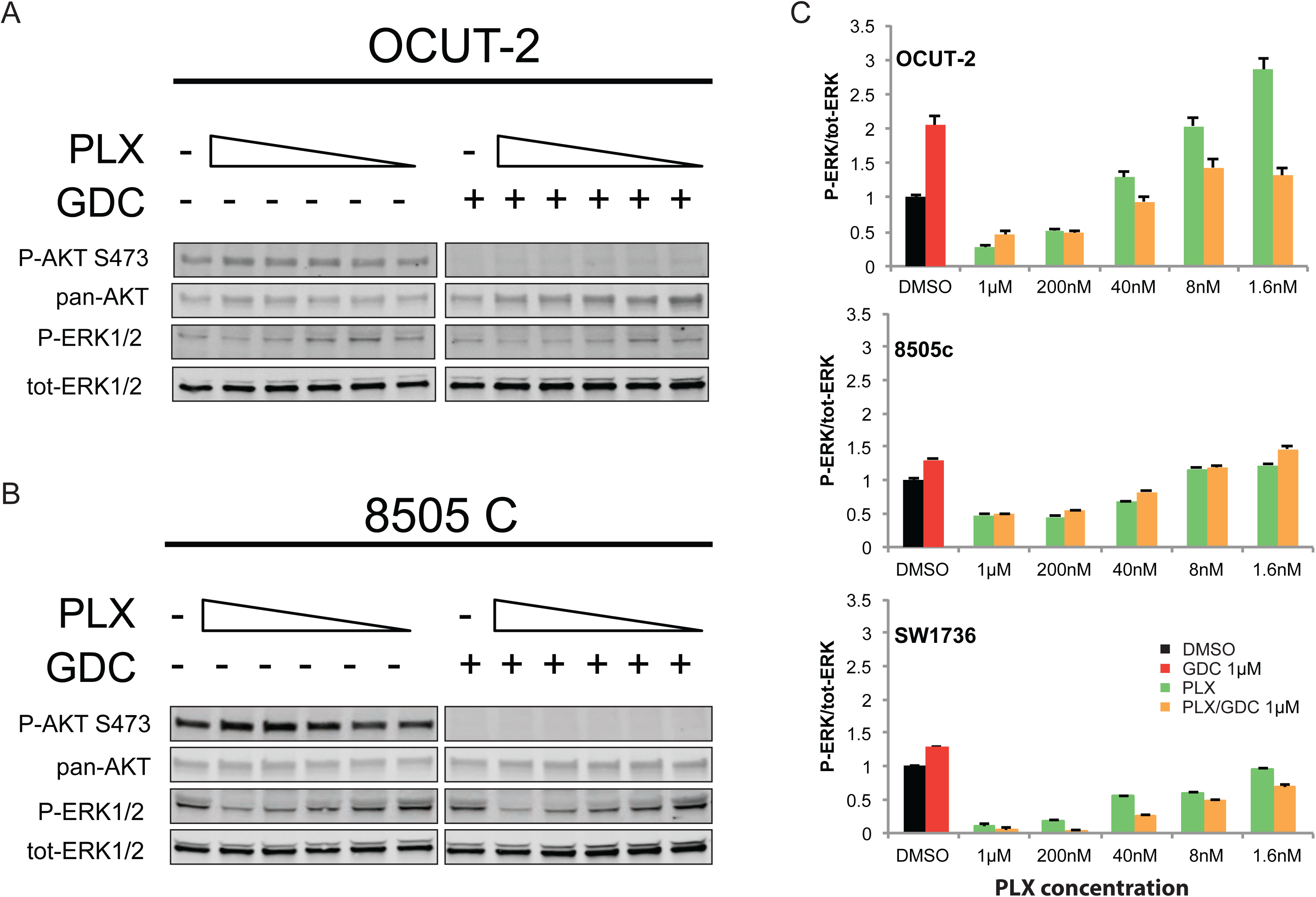
Paradoxical activation of ERK under BRAF^V600E^ specific inhibition treatment is also PI3K-dependent in human ATC cell lines. Representative western blots from total protein extracted from OCUT-2 (**A**) and 8505c (**B**) cells exposed to decreasing concentrations of PLX-4032 (from 1 μM to 1.6 nM) in presence or absence of GDC-0941 at 1 μM. All experiments were performed in triplicate and the quantifications are expressed as average ratios of the three independent experiments (**C**). The quantification of 8505C stands as a representative for both BRAF^V600E^ single mutant cell lines.

## Discussion

The BRAF^V600E^ mutation is detected in many cancer types such as melanoma, colon, ovarian, lung and prostate cancers [37,38]. In general it accounts for 8% of all mutations in human cancers [37]. The development of specific inhibitors against this mutation has raised a lot of interest in the clinics, for its potential lack of side effects. There is a strong rational to translate the success of vemurafenib from melanoma to other BRAF^V600E^ positive tumors like PTC or ATC. Unfortunately, case studies using vemurafenib in aggressive invasive-PTC and ATC show diverse results ranging from regression to no response [26–29].

We showed here that BRAF^V600E^ inhibition presented the expected response in BRAF^V600E^ single mutant mice (Fig. 1B). Tumor burden decreased at first then reached a plateau, reflecting the fact that after an initial response phase, MAPK signaling is transduced through BRAF^WT^ restoring a “normal” level of ERK phosphorylation (Fig 3B) resulting in no further clinical benefit of the drug. This was rather expected. On the other hand, when subjected to the same dose of PLX-4720, BRAF^V600E^ PIK3CA^H1047R^ double mutant tumors regrew after a very short initial response (Fig. 1C). This suggests that the additional PIK3CA^H1047R^ mutation resulted in a rapid resistance to BRAF^V600E^ specific inhibition. This was confirmed by the resistance break obtained by the inhibition of the PI3’K activity with the addition of GDC-0941 to the treatment regime. Consistently, recent studies in a model of colorectal cancer have shown that resistance to PLX-4720 treatment *in vivo* can be relieved by inhibition of PI3’K [39].

The initial phase of tumor reduction after PLX-4720 treatment in the double mutant model can be explained by tumor heterogeneity. Indeed, in this mouse model, tumors consist of rare normal follicles amongst large PTC and ATC foci (Fig 2A-E). Full Cre-driven recombination of both alleles from latent to mutant might not have occurred in every original cell. A portion of the tumors might be BRAF^V600E^-single mutant and still be responsive to PLX-4720. Interestingly, in the second double mutant cohort of mice, the resistance to PLX-4720 was already detectable after 10 days while GDC-0941 and drug combination already displayed a significantly tumor reduction effect (Fig. 4A) most likely due to a smaller proportion of single mutant cells left.

Even though treatment with GDC-0941 clearly induced a 20% reduction in the size of the tumor (Fig. 1C), possibly due to apoptosis elevation (Fig. 5), there was no clear beneficial effect in terms of histology (Fig. 2B). In terms of pathways activation, AKT phosphorylation level was reduced profoundly after 4 hours (Fig. 4B) but the effect seemed weaker after 10 days (Fig. 4B) and 10 weeks (Fig. 3A) of treatment. This suggests that there are also mechanisms of adaptation in this pathway, allowing restoration of AKT phosphorylation even when PI3’K is inhibited. The fact, that there is still a cytostatic effect when treating with GDC-0941, even though AKT phosphorylation level has partially recovered, demonstrates that AKT phosphorylation and PI3’K activity/mutation are not necessarily linked in this context [40]. This suggests that a mechanism downstream of PI3’K that does not necessary involve AKT could be responsible for tumor promotion. This is consistent with a recent publication from our group were PI3’K inhibition was combined with the MEK inhibitor PD-325901 [35].

Our most unexpected observation was that double mutant mice treated with PLX-4720 alone displayed a strong increase in ERK phosphorylation (Fig. 3A) after 10 weeks, that goes beyond recovery of ERK signaling. This was also observed 10 days and 4 hours after treatment (Fig. 4B). We could confirm this by immunofluorescence staining, where PLX-4720 treatment triggered an increase in the P-ERK signal in tumor cells (Fig. 4C). BRAF^V600E^ specific inhibition has been showed before to decrease P-ERK1/2 at first, then ERK activity will be restored by releasing the feedback inhibition loop between ERK and RAS, allowing “normal” RAF signaling through the wild-type allele of BRAF, as shown in our single mutant mice (Fig. 3B). The over-activation that we observe in the context of our PLX-4720 treated double mutant mice is however more reminiscent of the “paradoxical effect” of RAF inhibitors [41]. This paradoxical effect was found in melanoma patients treated with vemurafenib who developed squamous cell carcinoma [42] and explained by aberrant dimerization and enhanced signaling through BRAF^WT^ and CRAF. [43–46] Interestingly, this effect has been suggested to be used in used beneficially in wound healing [47].

The elevated ERK activation in the present study explained the worsening of the condition of the double mutant mice treated with PLX-4720 (Fig. 2E). Interestingly, in order to take place, paradoxical ERK activation requires a signal upstream activating CRAF. In this case, CRAF (also called Raf-1) could be activated by PI3’K [48]. Paradoxical ERK activation is most likely also the cause for the observed elevation of Ki67 in the tumors of PLX-4720 treated double mutant animals (Fig. 5A) as well as tumor progression (Fig. 2D&E). The elevation of the Ki67 index could also be caused by the observed progression of the tumors of PLX-4720 treated animals to ATC, since it has presents dramatically elevated proliferation index compared to well differentiated thyroid cancers. This progression to ATC was furthermore evidenced by the decreased TUNEL (Fig. 5B) and the decreased collagen deposition (Fig. 5C) since ATC are refractor to drug induced apoptosis. We could speculate that a paradoxical ERK activation might have driven tumor progression in the reported case of tumor progression observed with vemurafenib in clinical practice [27].

To assess whether this effect was transcriptionally regulated, we treated tumor bearing mice with one dose of PLX-4720 and dissected tumors 4 hours post treatment. The paradoxical effect could already be observed at that point (Fig. 4B/C) advocating for a non-transcriptionally regulated effect. Altogether this data suggests that paradoxical ERK activation is non-transcriptionally regulated and PI3’K dependent. Which is significantly different from the resistance observed in vivo by the Fagin group.

*In vitro*, the paradoxical effect was only observed in the OCUT-2 cell line (Fig. 6A) that has the same mutation pattern as the double mutant BRAF^V600E^/PIK3CA^H1047R^ mice but not in 8505c or SW1736 cells, (Fig. 6B) that have no reported alteration in PI3’K. These two cell lines represent the single mutant mice. The observed effect of paradoxical ERK activation in OCUT-2 cells could again be prevented when cells were concomitantly treated with GDC-0941 (Fig. 6A), once more showing that the paradoxical ERK hyper-activation is dependent on PI3’K activity. It is also important to note that the paradoxical effect is dependent on the drug concentration and might therefore have been missed in several studies using 1 μM PLX-4032. This can be explained by the fact that at this level PLX-4032 can inhibit all RAF isoforms and therefore also block CRAF, which is required for paradoxical activation. However, as this concentration of PLX-4032 is not applicable *in vivo*, we believe that our conditions are much more representative of physiological processes.

Drug combination treatment presented the strongest effect on tumor burden (Fig. 1C). Looking at TUNEL staining, GDC-0941 and drug combination treated animals could not be distinguished (Fig 5B). Apoptosis is a short-term process of programmed cell death. In general, cell debris disappears by extracellular enzyme digestion and leukocyte phagocytosis and is replaced by accumulation of fibrotic tissue, which is measured in daily clinical practice as response to chemotherapy in so-called regression scores in various human tumor types [49–51]. Deposition of fibrotic tissue was measured by Masson’s trichrome staining (Fig. 5C). Masson’s Trichrome staining showed significantly less deposition of fibrotic tissue in GDC-0941 treated animals when compared to combination treated animals. This revealed a slightly increased rate of cellular death in combination treated than in GDC-0941 treated animals that could only be visualized by a method that measures the accumulative effect of cell death. Whether this was due to apoptosis, necrosis or a different type of cell death remains unclear. With a similar Ki67 index, and TUNEL count, a further cell death mechanism might be involved under combination treatment (e.g. necrosis). It is also important to note that the areas that were mainly affected consisted of PTC, as demonstrated by the absence of PTC in tumors of combination treated animals and that ATC foci remained, meaning that this drug combination was mostly effective on PTC, but the effect on ATC remains elusive and would require more work *in vivo*.

Our data led us to propose the following mechanism for PLX-4720-induced ERK paradoxical activation and therefore resistance: BRAF^V600E^ specific inhibition leads to an increased formation of an active B/CRAF heterodimer and the complex is further stimulated by PI3’K [43–46,48,52] resulting in increased ERK phosphorylation as schematically presented in Fig.7. The two events, stabilization of a BRAF^V600E^/CRAF complex by BRAF^V600E^ specific inhibitors and activation of CRAF through PI3’K, are required to obtain paradoxically hyper-phosphorylated ERK1/2 that led to the worsening of the phenotype (Fig. 1). Our data are consistent with the fact that RAS activates PI3’K, and that newly arising skin tumors in vemurafenib treated BRAF^V600E^ mutant melanoma patients often carry activating mutations in RAS [42]. This novel *in vivo* mechanism of resistance clearly differs from published resistance mechanisms in melanoma, where resistance seems to be acquired during the course of treatment [20,21,23]. Thus, our data clearly improve the understanding of resistance mechanisms against vemurafenib.

**Figure 7:**
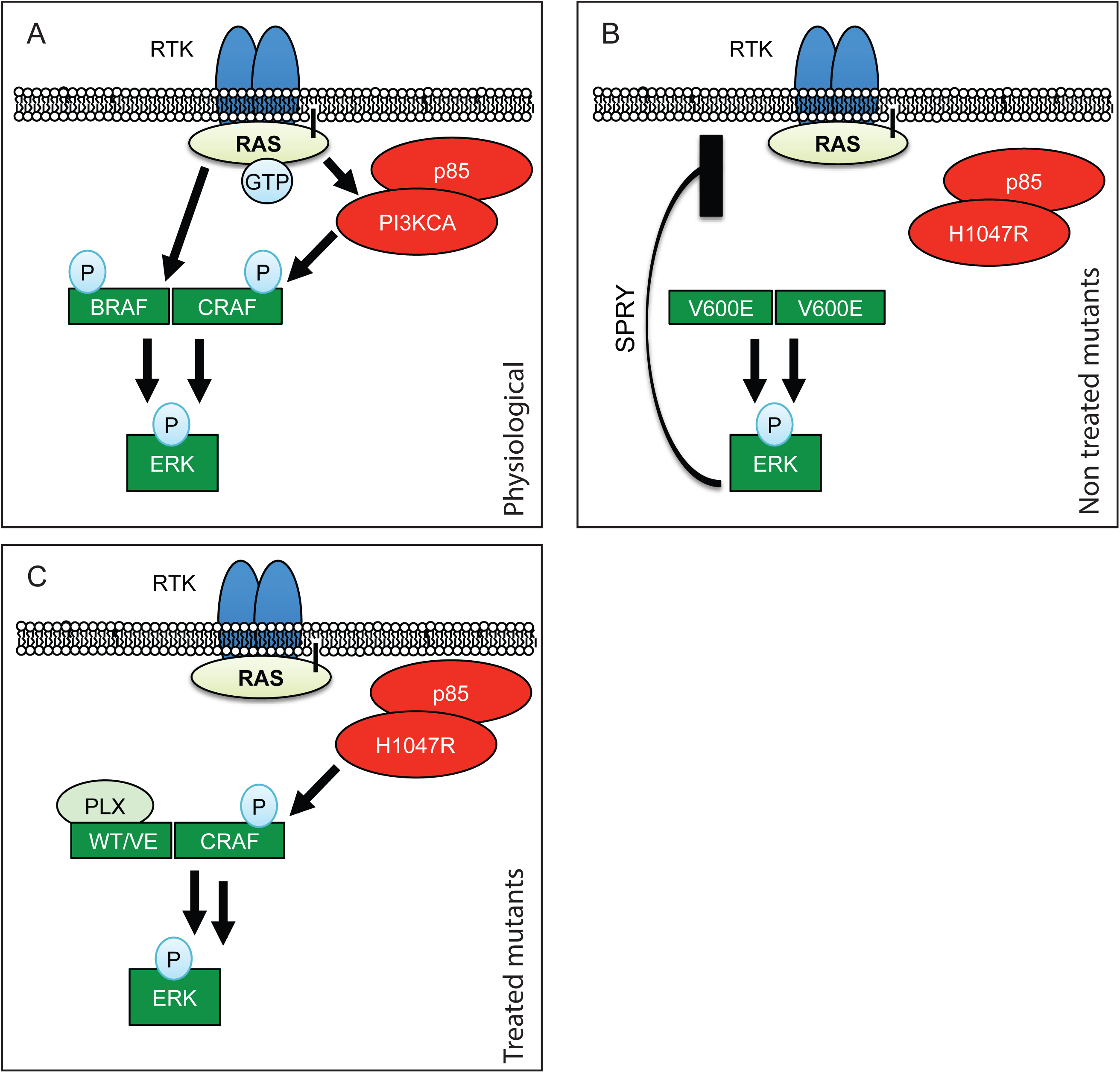
Schematic representation of RAF paradoxical activation. **(A)** In physiological conditions, upon Receptor Tyrosine Kinase (RTK) activation, receptor clustering induces RAS recruitment to the membrane. This results in dimerization of B/CRAF and subsequent phosphorylation that, *in fine,* induces ERK phosphorylation. **(B)** When BRAF is mutated to BRAF^V600E^ it is able to dimerize without RAS recruitment and induce ERK activation independently from upstream signals. ERK activation results in Sprouty’s up-regulation (SPRY) and therefore moderation of the pathway. **(C)** When treated with PLX-4720, BRAF^V600E^ is stabilized in a form that allows its dimerization with CRAF. This complex is then further stimulated by hyperactive PI3’K, leading to paradoxical ERK activation.

Finally, our findings suggest that selecting patients for targeted therapies based on their BRAF mutational status alone is not sufficient for choosing a successful therapy. The mutational status of *PIK3CA* should also be assessed, as treating BRAF^V600E^ positive patients carrying activating PI3’K mutations with BRAF^V600E^-specific inhibitors may lead to ERK hyper-activation and disease aggravation. Gene amplification measurements should also be considered, since more patients seem to be resistant to vemurafenib than suggested by the PI3’K activating mutation frequency. In conclusion, our data propose an explanation for the observed high rate of *a priori* drug-resistance to BRAF^V600E^ specific inhibitors seen in thyroid cancer. Nevertheless, we show that treating patients with RAF inhibitors like vemurafenib could be a valid approach in the appropriate clinical setting including prior mutational testing of the tumor and, if applicable, administered alone or combined with PI3’K inhibitors. This is also supported by the fact that even cell lines like 8505c, that show some resistance *in vitro* [53], are still responsive to PLX-4720 *in vivo* [54]. The combination of both PI3’K and BRAF^V600E^ inhibition may lead to a stronger response and should therefore be considered in the clinics, since a clear collaborative effect is visible *in vivo*. This drug combination or equivalent may be a good option for the treatment of aggressive inoperable PTC, but would require further investigation to clarify the effect *in vivo* on ATC.

## Material and Methods

### Cell lines

The BRAF^V600E^ mutant ATC cell lines 8505c and SW1736 were purchased at the Public Health England repository and cultured in RPMI-1640, 10% FCS, 2 mM L-glutamine, MEM NEAA (Thermo Fisher) 1:100 and P/S (100U Penicilin/ml and 0.1mg Streptomycin/ml). The ATC cell line OCUT-2 carries concomitant BRAF^V600E^ and PIK3CA^H1047R^ mutations [55]. It was kindly provided by Prof. James Fagin (Memorial Sloan Kettering Cancer Center), validated by our group by STR profiling (Microsynth, Switzerland), and cultured in DMEM medium 10% FCS, 2 mM L-glutamine, MEM NEAA (Thermo Fisher) 1:100 and P/S (100U Penicilin/ml and 0.1mg Streptomycin/ml). The mutational status for *PIK3CA* of all three cell lines was validated by sequencing. All cell lines were cultured for a maximum of 40 passages or 6 months; whichever limit was reached first.

### Drugs used

PLX-4720, PLX-4032 (vemurafenib), GDC-0941 and BKM-120 were purchased from Abmole Bioscience, Hong-Kong.

### Mice

All mouse experiments were performed in compliance with Swiss federal legislation and licensed by the Kanton of Bern. License Nr: BE120-13. Mice were kept in isolated ventilated cages, fed ad libitum in a 12/12 hours cycle of light and dark.

### Drug administration, mutations induction and tumor burden assay

*Braf^CA/+^; ThyroglobulinCre^ERT2^* single mutant or *Braf^CA/+^; Pik3ca^Lat/+^; ThyroglobulinCre^ERT2^* double mutant mice from a mixed FVB/C57BL6/F129 background were bred and mutations were induced by daily intraperitoneal injections of 1 mg tamoxifen diluted in peanut oil (100 μL) on five consecutive days. After two months of tumor growth, tumor bearing mice were treated by oral gavage with 30 mg/kg PLX-4720, 50 mg/kg GDC-0941 or the combination of both, formulated in a solution of 0.5% Hydroxypropylmethylcellulose (HPMC) (Sigma H7509) and 0.2% Tween 80 (Sigma P4780) six days per week. All mice weighed between 20 and 30 g at the start of the experiment. Group allocation for drug treatments was done randomly. Group size was determined using a power calculation assuming the following parameters: difference: 20%; variation: 20%; α 5%; β 50%. Beginning on the first day of drug treatment, tumors were measured by ultrasound measurement every week. For ultrasound measurements mice were anesthetized using 5 μl/g of body weight of a mixture of 0.1 mg/ml Dorbene, 0.5 mg/ml Dormicum and 5 μg/ml Fentanyl in 0.9% NaCl by intraperitoneal injection. The fur around the neck was epilated with Veet^®^ hair removal cream. Pictures were acquired with an ESAOTE MyLab Five ultrasound machine using a LA455 Probe (18 Mhz) from Siemens. After imaging, mice were taken out of anesthesia with 10 μl/g body weight of a mixture of 0.25 mg/ml Alzan, 5 μg/ml Flumazenil and 20 μg/ml Naloxon in 0.9% NaCl by subcutaneous injection. Images were analyzed using the ImageJ software. Evaluation of tumor burden was performed by th experimenter by measuring the surface of the biggest cross section in mm^2^ and normalized to the starting tumor burden of each mice for comparison.s

### Total protein preparation from cells

Cells were washed twice with cold PBS, recovered by scraping, then lysed in RIPA buffer (20 mM Tris pH 7.4, 150 mM NaCl, 1% Triton X-100, 0.1% SDS, 0.5% sodium deoxycholate) complemented with Halt(tm) Inhibitor Cocktail 1:100 (Pierce, ThermoFischier Scientific). The lysates were incubated 30 min on ice and then cleared by 15 min of centrifugation at 16,000 g at 4 °C. Protein concentrations were quantified by the BCA method (BCA Protein Assay Kit, Pierce, ThermoFischer Scientific). Lysates were prepared at 1-4 μg/μl in sample buffer (0.294 M sucrose, 2% SDS, 1 mM EDTA, 60 mM Tris pH 8.8, 0.05% Bromophenol blue, and 26 mM dithiothreitol) and the proteins analyzed by western blotting (see below).

### Total protein preparation from mice

Tissues were resected from anesthetized mice (10 mg/ml of Ketamin and 1.6 mg/ml Xylazin, at a dose of 10 μl/g body weight by intraperitoneal injection). After resection, the tissues were washed in ice-cold PBS (137 mM NaCl; 2.7 mM KCl; 18mM KH2PO4; 100mM Na2HPO4), snap frozen in liquid nitrogen and stored at −80°C until protein extraction. Proteins were extracted from tissue using RIPA buffer (see above). Halt(tm) Inhibitor Cocktail was added to the RIPA buffer to prevent dephosphorylation and protein degradation. 100 μl of RIPA plus Halt was added to the tissue samples for lysis using a QIAGEN TissueLyser LT at 50 Hz until complete homogenization (1-2 min). For large pieces of tissue, the amount of lysis buffer was increased. After complete tissue disruption, the sample was incubated on ice for 30 min and then centrifuged for 30 min at 4°C and 17,000 g. Protein concentration of the supernatant was evaluated as described above. All animals were euthanized 4 hours after their last drug administration.

### Western blotting

Protein extracts were either loaded separately (for quantifications) or pooled (for figure pictures) for each treatment condition. Proteins were run on TGX precast 4-20% gels (BioRad, Switzerland), transferred onto Trans-Blot transfer pack nitrocellulose (BioRad, Switzerland). Western blots were probed with the following antibodies and concentrations: Primary antibodies (dilution and catalogue number) from Cell Signaling (Purchased from Bioconcept AG, Switzerland): ERK1/2 (1:5000 9107); P-ERK1/2 (1:2000 4370); pan-AKT (1:2000 4691); pan-Akt (1:2000 6040); P-AKT-Ser473 (1:2000 4060) and actin (1:5000 or 1:10’000 A2066). Secondary antibodies, Li-Cor Biosciences (Bad Homburg Germany): IRDye 680RD Goat anti-Mouse IgG (H + L) (1:10’000, 926-68071), IRDye 800CW Goat anti-Rabbit IgG (H + L) (1:10’000, 926-32210). Blots were scanned using a Li-Cor ODYSSEY Sa fluorescent western blot scanner and quantified with the ODYSSEY Image Studio software.

### Immunofluorescence

Tissue samples for histology were recovered concomitantly to those recovered for proteins, washed in cold PBS and fixed overnight in a neutrally buffered 10% formalin solution (Sigma HT501128). Paraffin-embedded tissue was sectioned to 5 μm. The sections were rehydrated after paraffin removal and targets were retrieved in Tris (Sigma T1503) 10 mM, EGTA (Sigma E4378) 0.5 mM pH=8.0 solution. The sections were blocked three times for 10 minutes in a buffer of 1% BSA, 0.2% gelatin, and 0.05% Saponin (Sigma 47036) in PBS. The primary antibody KI67 (1:300; Abcam, ab16667) was diluted in 0.1% BSA and 0.3% Triton X-100 in PBS. Other primary antibodies were Ck-19 (1:300; DSHB, 5605s), galectin 3 (1:300 Abcam ab53082), vimentin (1:300 Bioconcept 5741S) and P-ERK1/2 (1:300 Bioconcept 4370L) The primary antibody was incubated overnight at 4 °C. The slides were then washed three times with 0.1% BSA, 0.2% gelatin and 0.05% Saponin in PBS. The secondary antibody was goat anti-rabbit 488 (Life Technologies A-11034 1:500) complemented by DAPI (Sigma 32670) at 5 μg/ml to counterstain nuclei and incubated for 1 hour at room temperature. Slides were scanned with a Panoramic Midi Scanner (Sysmex/3DHISTECH Switzerland/Hungary). Analysis and quantification was performed on whole tumor sections with the QuantCenter software (Sysmex/3DHISTECH Switzerland/Hungary).

### Hematoxylin-Eosin Staining

For histological analysis, tissue samples were processed as described above and stained with Hematoxylin (Sigma GHS132) and Eosin (HT110132) following standard protocols.

### TUNEL

For TUNEL staining tissue was processed and sectioned according to the procedure described in the preparation of Immunofluorescence slides. Rehydration was also done as described in the Immunofluorescence section. Sections were pretreated with proteinase K (Sigma P2308) with an activity of 0.6 units/ml for 10-20 min in a humidifying chamber at 37°C before they were left to cool at room temperature for 10 min. Subsequently, sections were washed twice in PBS 1x Tween20 (Sigma P9416) 0.1% for 2 min each. To decrease background signal, endogenous biotin was blocked using a kit from Thermo Fisher (E21390). Then sections were pre-incubated in TdT reaction buffer containing 25 mM TRIS-HCl pH 6.6 (Sigma T1503), 200 nM Sodium Cacodylate (Sigma C0250), 0.25 mg/ml BSA and 1 mM Cobalt Chloride (Sigma C8661) for 10 minutes at room temperature. After pre-incubation, the sections were incubated in TdT reaction mixture for 1-2 h at 37°C in a humidifying chamber. TdT reaction mixture contains Terminal deoxynucleutidyl Transferase (TdT) (Sigma 3333566001) and Biotin 16-dUTP (Roche 11093070910). The reaction was stopped by incubating the sections with a buffer containing 300 mM NaCl (Sigma S9888) and 30 mM sodium citrate (Sigma S4641) for 10 min at room temperature. The sections were then washed three times for 2 min in PBS containing 0.1% Tween20. Before labeling, the slides were blocked with a solution of 3% BSA (Sigma A7906) in PBS 1x, followed by three washing steps (2 min each) with PBS + Tween20 (0.1%). For detection, slides were incubated with a solution containing streptavidin-Alexa Fluor 633 (1:2000 S21375 from LuBioScience) and DAPI 5 μg/ml (32670 Sigma Aldrich). Slides were rinsed with PBS 1x and coverslips were mounted using anti-fading fluorescent mounting medium from DAKO (S3023).

### Masson’s Trichrome

For Masson’s Trichrome staining, tissue was processed and sectioned according to the procedure described in the preparation of Immunofluorescence slides. Rehydration was also done as described in the Immunofluorescence section. Following rehydration slides were incubated with Bouins fixative solution containing 75% saturated picric acid (Sigma P6744), 25% of 37% formaldehyde solution (Sigma 252549) and 5% acetic acid (Sigma 33209) at room temperature overnight. After fixation nuclei were stained with Weigert’s hematoxylin solution containing 0.5% hematoxylin (Sigma H9627), 0.5% concentrated HCl (Sigma 320331), 2% of a 29% ferric chloride solution (diluted from 45%) (Sigma 12322), and 47.5% (diluted from 100%) ethanol (Sigma 02860) in distilled water for 10 min at room temperature. Slides were then rinsed under warm running tap water for 10 min, before they were rinsed in distilled water. The extra-nuclear tissue was then stained with the Biebrich Scarlet-Acid Fuchsin solution containing 89% of a 1% Biebrich Scarlet solution (Sigma B6008), 10% of a 1% Acid Fuchsin (Sigma F8129) solution and 1% of acetic acid for 10 min at room temperature, followed by rinsing in distilled water until clear. To remove the stain from the collagen for differentiation, the slides were subsequently incubated with a solution containing 50% of a 5% phosphomolybdic acid solution (Sigma HT153) and 50% of a 5% phosphotungstic acid solution (Sigma HT152) for 10 min at room temperature. Then slides were transferred to a solution containing 2.5% Aniline Blue (Sigma 415049) and 2% acetic acid in distilled water for 10 min at room temperature for counterstaining of collagen. Slides were then dehydrated rapidly in a succession of two baths of 1% acetic acid followed by 1 bath of 95% ethanol and two baths of 100% ethanol. Slides were then left in a xylene bath until mounting in Eukitt^®^ Quick-hardening mounting medium (Sigma 03989).

### Statistical methods

All statistical analyses were performed in GraphPad®. Tumor growth experiments were analyzed using the Mann-Whitney test to compare the groups. The remaining experiments were analyzed using one-way ANOVA with a Fisher’s exact test for post hoc testing. In cases where there were only two groups to compare, a two-tailed t-Test was used. Significance is displayed using stars with one star meaning a resulting p-value of smaller than or equal to 0.05. Two stars means a p-value of smaller than or equal to 0.01. Three stars stands for a p-value of smaller than or equal to 0.001. If a p-value was greater than 0.05 the difference between the two concerned groups was regarded as not significant.

## Ethics Approval

Mice were kept, treated and euthanized according to the Swiss federal guidelines. The experimental protocol was approved by the Bernese cantonal ethical commission for animal experimentation (License number: BE120/13).

## Author Contributions

***In vivo* experiments:** MAR and OEM.

***In vitro* experiments**: DRD.

**Tissue staining:** MAR and AS.

**Wester slots:** MAR.

**Data analysis:** MAR:

**Animal breeding, genotyping:** AS and MAR.

**Histopathological analysis:** MSD.

**Manuscript writing:** RPC, MAR, DRD, MSD and WAP.

**Figure preparation:** MAR, DRD, RPC.

**Design:** RPC.

**Study supervision:** RPC.

## Acknowlegements

Special thanks to Prof. Martin McMahon for his former support, mentoring and for the *Braf*^CA^ mice. Further, special thanks to Prof. Engelhard, Dr. Deutsch and Dr. Benarafa for allowing us to keep our mice in the vivarium at the Theodor Kocher Institute. Finally, to Prof. Dimitrios Fotiadis for his pushes in the right direction. Also, we would like to acknowledge the Microscopy Imaging center (MIC) of the University of Bern. Mr Roelli and Mr ElMokh are enrolled in the Graduate School for Cellular and Biomedical Sciences (GCB) of the University of Bern, that we would like to acknowledge for the training provided.

---

*Supplementary figure 1:* Paradoxical activation of ERK abrogated by BKM-120.

Representative western blot from total protein extracted from OCUT-2 **(A)** and 8505c **(B)** cells exposed to decreasing concentrations of PLX-4032 (from 1μM to 1.6 nM) in presence or absence of BKM-120 at 1 μM. All experiments were performed in triplicate. The quantifications are expressed as average ratios of the three independent experiments.

